# FungalRoot: Global online database of plant mycorrhizal associations

**DOI:** 10.1101/717488

**Authors:** Nadejda A. Soudzilovskaia, Stijn Vaessen, Milargos Barcelo, Jinhong He, Saleh Rahimlou, Kessy Abarenkov, Mark C. Brundrett, Sofia Gomes, Vincent Merckx, Leho Tedersoo

**Author notes:** corresponding authors with equal contribution. The data inquiries should be addressed to Nadejda Soudzilovskaia. authors equally contributed to the paper, being the main data compilers (Stijn Vaessen, Milargos Barcelo).

## Abstract

- The urgent need to better understand profound impacts of mycorrhizas on functioning of terrestrial ecosystems, along with recent debates on resolving plant mycorrhizal associations, indicate that there is a great need for a comprehensive data of plant mycorrhizal associations able to support testing of ecological, biogeographic and phylogenetic hypotheses.
- Here present a database, FungalRoot, which summarizes publicly available data on plant mycorrhizal type and intensity of root colonization by mycorrhizal fungi, accompanied by rich meta-data. We collected and digitized data on plant mycorrhizal colonization intensity published until April 2019 in 9 globally most important languages. The data were assessed for quality and updated for plant taxonomy.
- The FungalRoot database contains 36,303 species by site observations for 14,870 plant species, tripling the previously available amount in any compilation. The great majority of ectomycorrhizal and ericod mycorrhizal plants are trees and shrubs, 92% and 85% respectively. The majority of arbuscular mycorrhizal and of non-mycorrhizal plant species are herbaceous (50% and 70%).
- Besides acting as a compilation of referenced observations, our publicly available database provides a recommendation list of plant mycorrhizal status for ecological and evolutionary analyses to promote research on the links between above- and belowground biodiversity and functioning of terrestrial ecosystems.

## Introduction

Mycorrhizal interactions with fungi represent one of the key innovations of terrestrial plants. Mycorrhiza is a mutualistic association between plant roots and fungi, where plants provide photosynthetically derived carbohydrates to fungi and fungi deliver nutrients and water to plants and offer protection from abiotic and biotic stress (Smith & Read, 2008). Based on tomy and phylogeny, four principal types of mycorrhiza are recognized: arbuscular mycorrhiza (AM), ectomycorrhizal (EcM), ericoid mycorrhiza (ErM) and orchid mycorrhiza (OM) (Brundrett & Tedersoo, 2018). While most vascular plant species form mycorrhizal symbiosis of only one type, AM-colonized plants may sometimes co-occur with EcM and ErM fungi (Smith & Read, 2008; Brundrett & Tedersoo, 2018).

Depending on mycorrhizal types and particular species, mycorrhizal fungi may build extensive mycelial networks that sustain nutrient acquisition and promote plant seedling establishment (Leake *et al.*, 2004; Soudzilovskaia, N.A. *et al.*, 2015). Mycorrhizal types differ in plant nutrition and therefore affect plant carbon allocation strategies (Veresoglou *et al.*, 2012b), litter quality (Cornelissen *et al.*, 2001) cf (Koele *et al.*, 2012) and decomposition (Cornelissen *et al.*, 2001; Koele *et al.*, 2012; Elumeeva *et al.*, 2018), biogeochemical cycles (Veresoglou *et al.*, 2012a; Soudzilovskaia *et al.*, 2015; Averill & Hawkes, 2016; Tedersoo & Bahram, 2019), and plant community composition (van der Heijden *et al.*, 1998; Klironomos *et al.*, 2000; Klironomos *et al.*, 2011; Elumeeva *et al.*, 2018). Information about mycorrhizal type and colonization intensity of mycorrhizal infection of plant roots is crucial for understanding plant and fungal effects on ecosystem-level and global biogeochemical processes (Phillips *et al.*, 2013; Terrer *et al.*, 2016).

Plants also differ in the level of root colonisation by mycorrhizal fungi, which may have an effect on the efficiency of nutrition (Karst *et al.*, 2008; Hoeksema *et al.*, 2010; Treseder, 2013) or protection against pathogens (Smith & Read, 2008). Much of the colonisation level seems to be related to plant and fungal identity but also to seasonality and environmental conditions (Klironomos, 2000; Maltz & Treseder, 2015; Hoeksema *et al.*, 2018). Further, the data about root colonization by mycorrhizal fungi provides insights into the level of intimacy of the plant-fungal relation, linked to the plant nutrition effectiveness and plant global environmental drivers (Soudzilovskaia *et al.*, 2015). Several plant species so-called ‘facultatively mycorrhizal plants’ may develop mycorrhizas in certain conditions but remain non-mycorrhizal in other conditions, depending on nutrient availability and neighbouring plants (Brundrett, 2009)

However, the type and level of infection by mycorrhizal fungi is unknown for the great majority of vascular plants and, when available, this information is scattered in multiple narrow-focused data sets, most of which cover specific Earth regions or mycorrhizal types. Many sources of mycorrhizal records contain multiple errors, which have accumulated and passed on through literature reviews. Many of these errors are derived from alternative interpretations of mycorrhiza and mycorrhizal types, which is especially common in old literature (Brundrett & Tedersoo, 2019; Bueno, 2019). Unfortunately, these incorrect observations have been commonly used in traits-based case studies or meta-analyses without critical evaluation of the source reliability, which may have caused slight to fatal errors in interpretation and conclusions literature (Brundrett & Tedersoo, 2019). Furthermore, most of data compilations lack geographical information and any environmental metadata about the study sites. Also, substantial part of fundamental mycorrhizal research has been published in languages other than English or German or French, which have remained mostly overlooked in reviews and data sets. Finally, virtually none of the existing data compilations distinguish between research focused on all mycorrhizal types detected for a particular species and specific mycorrhizal types (ignoring others when present).

Here, we present a global database FungalRoot, which accumulates information about plant mycorrhizal status and root colonization intensity, The FungalRoot database was assembled based on previously published reviews, local databases and a large number of yet neglected case studies and recent studies published in nine globally most important languages. The database enables to distinguish between reports of a presence of a particular mycorrhizal type, and reports where the plants were checked for *all* existing mycorrhizal types. In addition, our database provides information about the locality, ecosystem type, soil chemical data, and the method of mycorrhizal assessment that enable users to build more specific, local reference databases. FungalRoot offers possibilities to provide curator and third-party expert opinion regarding data reliability. Based on the current version of the database we provide a genus-level recommendation list for mycorrhizal type assignment of vascular plants, considering taxonomic information, habitat and phylogenetic conservation (Brundrett, M & Tedersoo, L, 2018). This list is also included into the FungalRoots database as a separate and, as well is open for third-party expert opinion. This data considerably advances the previously available major check list of plant mycorrhizal status (Wang & Qiu, 2006; Akhmetzhanova *et al.*, 2012) in number of plant taxa considered, and in the accuracy of mycorrhizal type diagnoses. The genus-level recommendation list for using the mycorrhizal trait in comparative studies and meta-analyses, in which mycorrhizal types are not empirically determined.

## Methods

### Literature compilation

We combined data from 1775 sources of literature including articles obtained through Google Scholar searches, large compilations of information on mycorrhizal colonization type and intensity in plants (Harley & Harley, 1987; Wang & Qiu, 2006; Akhmetzhanova *et al.*, 2012; Hempel *et al.*, 2013; Soudzilovskaia, *et al.*, 2015; Gerz *et al.*, 2016), and authors’ personal literature collections. For the Google Scholar search, we used boolean search ‘mycorrhiza’ AND ‘colonisation’ AND ‘name of each country’ in English and in other major languages (incl. German, Chinese, Farsi, French, Indonesian, Portuguese, Russian, Spanish). The sources were downloaded from the Internet, acquired from the authors or ordered using interlibrary loans. We focused mainly on papers with observations on at least five species or >10 observations for a lower number of species separable by space or specific treatments. Large data compilations were traced to the original references in order to add geographical and ecological metadata, check for methods and avoid accumulating errors.

Presence of mycorrhizal status information of plant species or genus was the minimum requirement to include observations in the database. In cases when the data on root colonization intensity by specific mycorrhizal type was reported, we included this data as well along with the method of colonization assessment. All collected references were carefully checked for information about geographical location, environmental and habitat conditions (See Table 1 for the lists of included variables and character states).

**Table 1.** Description of FungalRoot database fields.

| **Publication** | *Field explanation* |
| --- | --- |
| *dataset ID* | The unique number identifying the observation |
| *publication doi* | DOI number of the original reference (“unpublished”, when the reference is not published) |
| *original reference* | Name of original references from which the records were extracted (in APA format) |
| *non original reference* | In the original is unchecked, the indirect reference (in APA format) |
| *original_ref_checked [y/n]* | For compilations, this field specified if the original reference was checked or not |
| *chinese name* | In case of Chinese publications, the original study title |
| **Observation location** |  |
| *plot_name* | Name of the plot the sample belongs to according to the original publication |
| *habitat_naturality* | Habitat of plants (selection: natural, plantation, nursery, greenhouse, pot, axenic, wasteland, early successional, other) |
| *biome* | Specific biome where the observation was made. Selection: anthropogenic cropland; anthropogenic rangeland; anthropogenic urban; desert temperate; desert tropical; forest Mediterranean; forest subpolar coniferous; forest temperate broadleaf; forest temperate coniferous; forest tropical broadleaf; forest tropical coniferous; grassland flooded; grassland temperate; grassland tropical; mangrove; tundra; aquatic: freshwater lake; aquatic: freshwater river; aquatic: marginal sea; aquatic: marginal salt marsh; other |
| *Country* | Country where the observation was made. Selection form a list. |
| *Collection date* | Date of sampling (YYYY-MM-DD; YYYY-MM, YYYY or XXXX- |
|  | MM-DD if year is unknown) |
| *Latitude* | Latitude of the sampling location (WGS84) |
| Longitude | Longitude of the sampling location (WGS84) |
| Elevation | Elevation of the sampling site (m above sea level) |
| **Environmental metadata** |  |
| pH | Soil pH value; provided in a case of single measurement |
| pH_min | Minimum value of soil pH |
| pH_max | Maximum value of soil pH |
| pH_range | Value range of soil pH |
| pH method | Method used for determining soil pH. Selection: 'water' or 'KCl' |
| Total organic carbon | Total organic carbon content of the soil (g C/kg soil). When original values were expressed in g Org Matter/ kg soil, a conversion of 0.48g Org Matter = 1g C was applied |
| C_min | Minimum value of soil organic carbon concentration (g C/kg soil) |
| C_max | Maximum value of soil organic carbon concentration (g C/kg soil) |
| C_range | Value range of soil organic carbon concentration (g C/kg soil) |
| C_unit | The unit used for soil organic carbon concentration (g C/kg soil) |
| Total organic carbon method | Method used for determining the total soil organic carbon. Selection: Kjeldahl; elemental analyser; furnace |
| Total nitrogen | Total nitrogen content of the soil sample (g N/kg soil). Nitrate and ammonium values are calculated based on atom mass, coefficients 0.226 and 0.778, respectively) |
| N_min | Minimum value of total soil nitrogen concentration (g N/kg soil) |
| N_max | Maximum value of total soil nitrogen concentration (g N/kg soil) |
| N_range | Value range of total soil nitrogen concentration (g N/kg soil) |
| Total nitrogen method | Method used for determining the total soil nitrogen. Selection: Kjeldahl; elemental analyser |
| Total phosphorus | Total phosphorus concentration in the soil sample (mg P/ kg soil) |
| Available phosphorus_mean | Available phosphorus concentration in the soil sample (mean value) |
| P_min | Minimum value of available phosphorus concentration |
| P_max | Maximum value of available phosphorus concentration |
| P_range | Value range of available phosphorus concentration |
| P_unit | The unit used for available phosphorus concentration |
| Available phosphorus method: | Reference or method used for determining the concentration of soil phosphorus. Selection: oxalate; ammonium acetate; Bray; water; Olsen; other |
| Potassium | Total concentration of soil potassium |
| K_min | Minimum value of potassium concentration |
| K_max | Maximum value of potassium concentration |
| K_range | Value range of potassium concentration |
| K_unit | The unit used for potassium concentration |
| Calcium | Total concentration of calcium |
| Ca_min | Minimum value of calcium concentration |
| Ca_max | Maximum value of calcium concentration |
| Ca_range | Value range of calcium concentration |
| Ca_unit | The unit used for calcium concentration |
| Magnesium | Total concentration of magnesium |
| Mg_min | Minimum value of magnesium concentration |
| Mg_max | Maximum value of magnesium concentration |
| Mg_range | Value range of magnesium concentration |
| Mg_unit | The unit used for magnesium concentration |
| **Host plant description** |  |
| Species | Genus name and epithet of original observations. When epithet is not given, only genus name was is recorded. The recorded observations were checked against the Plant List database (www.plantlist.com) |
| Number of plants | N, the number of individuals of the plant species studied |
| *Host age* | Age of the host plant observed. Selection: <1 months;1 months (30-60 d); 2 months (61-90 days); 3 months (91-120 days); 4 months (121-150 days); 5 months (151-180 days); 6 months (181-210 days); 7 months (211-240 days); 8 months (241-270 days) 9 months (271-300 days); 10 months (301-330 days); 11 months (331-365 days); 12 months (365-384 days); <1 year (30-365 days); 12-24 months (365-730 days); 2-10 years (sapling); >10 years |
| **Plant mycorrhizal colonization** |  |
| *mycorrhiza_type* | Mycorrhizal type detected. Selection: AM (no others); AM (others not addressed); EcM (no others); EcM (others not addressed); ErM; OM; non-mycorrhizal (checked for all types); non-EcM (others not addressed); non-AM (others not addressed); EcM-AM; ErM-AM; ErM-EcM; AM-like (non-vascular plants); EcM-like (non-vascular plants); ErM-like (non-vascular plants); OM-like (non-vascular plants) |
| *AM intensity* | Extent of root system colonized by AM fungi |
| *AM frequency* | Percentage of individual plants colonized by AM fungi |
| *AM method* | Method used to determine AM fungi colonization intensity. Selection: McGonigle et al. 1990: RLC (%); Phillips & Hayman 1970: RLC (%); Selivanov 1981: RLC (%); scale 0-5 (Kormanik & McGraw 1982); scale 0-4 (Peuss 1958/Mejstrik); scale 0-3; Giovannetti & Mosse 1980: slide-length; Giovannetti & Mosse 1980: gridline intersect; Herper 1977: colorimetric; Sieverding,1991: RLE; qPCR; molecular identification; simple observation; synthesis; other (% scale); other |
| *EcM intensity* | Extent of root system colonized by EcM fungi |
| *EcM frequency* | Percentage of individual plants colonized by EcM fungi. |
| *EcM method* | Method used to determine EcM colonization intensity. Selection: root tips colonized (%); root length colonized (%); scale 0-4; scale 0-3; scale 0-2; qPCR; molecular identification; EcM tips/m root; simple observation; other |
| *ErM intensity* | Extent of root system colonized with ErM fungi |
| *ErM frequency* | Percentage of individual plants colonized with ErM fungi. Relevant when intensity is not given. |
| *ErM method* | Method used to determine ErM fungi colonization intensity. Selection: root length colonized (%); scale 0-4; scale 0-3; qPCR; molecular identification; simple observation; other |
| *OM intensity* | Extent of root system colonized with OM fungi |
| *OM frequency* | Percentage of individual plants colonized by OM fungi. Relevant when intensity is not given. |
| *OM method* | Method used to determine OM colonization intensity. Selection: root length colonized (%); scale 0-4; scale 0-3; scale 0-2; qPCR; molecular identification; simple observation; other. |
| *remark:*<br>*Mycorrhiza type* | A note on reported mycorrhiza type or determination method. |
| *Curator remark*<br>*1: name* | Name of the expert providing opinion |
| *Curator remark*<br>*1: comment* | Comment |
| *Curator remark*<br>*2: name* | Name of the second expert providing opinion |
| *Curator remark*<br>*2: comment* | Comment |

Data about site soil conditions were added to each record when available. Nitrate (NO^−3^) or ammonium (NH^+4^) values were converted to N based on atomic mass. Eg. X mg NO-3 /kg = X * 14/62 mg N /kg, as the atomic mass ratio between N and NO-3 is 14/62. Similarly for NH^+4^ with atomic mass ratio of 14/18 between N and NH^+4^. Minimum and maximum values, or ranges, were added when available.

### Assessment of mycorrhizal types

Here we follow the mycorrhiza definitions of Brundrett & Tedersoo (2018) that were largely built on Smith & Read (2008). Because the associated fungi were rarely identified and their benefits to plants were not addressed in studies addressing mycorrhizal status or level of colonisation of natural plants, we relied solely on the morphological criteria in most cases (except Australian studies in 1980s and early 1990s that involved synthesis experiments). In brief, the presence of intracellular arbuscules, coils or pelotons was required to consider plants AM, ErM or OM, respectively. For EcM, the presence of a Hartig net or a well-developed mantle (>1 hyphal layer) was required. Although Bueno (2019) argued that arbuscular colonisation is not required for functional AM symbiosis, there is ample evidence that well-colonized plants perform better in terms of nutrition and protection from pathogens and that incapacity of forming arbuscules is a characteristic of NM or facultatively mycorrhizal plants in natural conditions. Therefore, we considered only hyphal root colonisation and molecular identification from root samples insufficient to consider the plants mycorrhizal. Hyphae of AM, EcM and ErM fungi proliferate in surface-degraded roots of plants belonging to other mycorrhizal types (Toju *et al.*, 2014).

Misdiagnoses of mycorrhizal types are a common problem in scientific literature (Brundrett, 2017; Tedersoo & Brundrett, 2017; Brundrett & Tedersoo, 2019) and these could lead to errors in analyses based on data collected by literature compilations (Brundrett & Tedersoo, 2019). We considered it important to report in the database the original diagnosis of mycorrhizal type provided by in the original publication. Simultaneously, we examined each record in our database against contemporary knowledge of plant species mycorrhizal types (consensus of records in this study; specific information in (Merckx, 2013; Kohout, 2017; Tedersoo & Brundrett, 2017; Brundrett & Tedersoo, L, 2018; Brundrett & Tedersoo, 2019). Based on this examination, we provided one or two expert opinions commenting on the reliability of the original diagnosis for each contradictory record (see subsection “Data records”). Based on the database records and the expert opinions, we prepared a recommendation list of mycorrhizal status at the plant genus level (Table S1). Here we considered individual studies of low reliability and excluded these from further comparisons if >20% of records therein conflicted with other studies. We anticipate, however, that differences in NM and AM habit may exist especially in facultatively mycorrhizal plants and seasonally, depending also on age and environmental conditions (Bueno, 2019).

Based on individual reports for species, we assigned mycorrhizal type or NM status to the entire genus if >67% of reports (at least two observations) converged. In putatively AM and NM groups, there were multiple genera that were reported to be either AM or NM in 33%-67% of occasions. These facultatively arbuscular mycorrhizal taxa were encoded as AM-NM. In predominately AM and EcM plant families, we considered a single positive report sufficient to consider the genus mycorrhizal. If there was a single NM report in these mycorrhizal families, the mycorrhizal status was considered unsettled, to avoid considering these prematurely non-mutualistic or the report as unreliable. For genera that had no reports or single negative reports or two conflicting reports, we further relied on the list of putative NM plant families (Brundrett & Tedersoo, 2018) and EcM plant genera (Tedersoo, 2017) and studies considered unreliable in the first phase. Several aquatic and heterotrophic plant genera in the putatively NM plant families were considered as AM-NM because of multiple independent evidence for arbuscule formation.

According to our data, 86 plant families lack information about mycorrhizal types (Table S2). Following (Brundrett & Tedersoo, 2018), we considered that *Brassicales*, *Caryophyllales* and *Cyperales* included multiple families with mostly NM or AM-NM species. In *Brassicales*, we partly relied on the distribution of mycorrhiza-related genes (Delaux et al. 2014). If these reports were missing (*Brassicales*) and for other putatively AM-NM orders, we considered the mycorrhizal status of unstudied families unsettled. For AM orders that contain only AM genera, we considered these families as putatively AM. We also included specific comments on species of EcM plants that have a larger group of congeneric AM species (*Pisonia, Persicaria, Kobresia*). For Australian Fabaceae, Goodeniaceae, Myrtaceae and Asteraceae, this was unfeasible because of paucity of such information. We identified only a single consistently NM plant species *Astragalus alpinus* within a mycorrhizal genus.

### Technical validation

For correction and standardization of the species names included in the database, all observations were checked using the Taxonomic Name Resolution Service (TNRS) (http://tnrs.iplantcollaborative.org/). When partial matches were found, species names were corrected manually according to best suggestion given by the TNRS. When no suggestions were given, the species name was checked in the original papers. If after this step the species name could not be corroborated, the record was removed, or, where possible, treated at the level of genus.

### Data Records

The data are organized into five categories: (1) observation identification; (2) location; (3) soil conditions; (4) host plant description; and (5) description of mycorrhizal colonization (Table 1). The fields for literature references refer to one particular study and include ‘publication_doi’ (for a Digital Object Identifier, DOI, of the citation) and ‘original_reference’ (full text citation in GoogleScholar APA format, necessary for older literature with no DOI or other alphabet). Chinese, Japanese, Persian, Arabic, Cyrillic, etc. alphabets also conform to this field, although sources in these languages except Chinese have been converted or translated to the main format during data management or within previous reviews for simplicity. The field ‘plot_name’ enables to segregate the study into smaller units by location but also by time, treatment or any custom difference. It is represented by the name of locality or locality-by-treatment combination. All records within a plot have the same geographical coordinates. Identical plot names in different studies are not matched unless their coordinates match.

For the location category, ‘habitat naturality’ enables selection amongst eight possibilities (plus ‘other’ if none conform) that are related to the experimental conditions or habitat structure; ‘biome’ provides information about the overall climate and vegetation type; ‘country’ represents a user-selected field for countries (autonomous and overseas regions separately) following MIMARKS standards; ‘latitude’ and ‘longitude’ represent geographical coordinates, whereas ‘elevation’ represents altitude; ‘collection_date’ depicts date of observation

To enable in-depth meta-analyses, we have included 12 fields for soil chemical parameters that are most commonly reported in mycorrhiza literature, along with the description of methods for their assessments. The fields ‘pH’, ‘pH_min’, ‘pH_max’, ‘pH_range’ and ‘pH_method’ denote the value and measurement method for determining soil pH. The fields ‘total_organic_carbon’ and ‘total_organic_carbon_method’ are used for stating the value (g/kg soil) and determination method for soil organic carbon content. Similarly, ‘total_nitrogen’ and ‘total_nitrogen_method’ indicate the value (g/kg) and method of determination for total soil nitrogen. The fields ‘total_phosphorus’, ‘total_available_phosphorus’ and ‘total_available_phosphorus_method’ indicate concentration of total phosphorus (mg/kg soil; method, destruction) or available phosphorus (mg/kg soil) and its method of measurement. ‘Potassium’, ‘calcium’ and ‘magnesium’ represent fields for K, Ca and Mg concentrations (mg/kg soil; method, destruction).

There are three fields for plant species. One of the most important fields is ‘species’, which represents the taxon studied. If no epithet is given, the taxon is identified to genus level. The field ‘number_of_individuals_studied’ represents the sample size of the original study. The ‘host_age’ represents a selectable field of the age of particular individuals, ranging from <1 month to >10 years.

Information about mycorrhizal type and colonisation intensity and frequency are given in a suite of 13 fields due to data complexity. The field ‘mycorrhiza_type’ is a selection of 15 options indicating combinations from single mycorrhiza types to dual mycorrhizal colonisation and specifying whether other types were addressed or not. We find these possibilities important to be considered in scientific analyses, as they allow distinguishing between negative reports that may be derived from the lack of survey for other mycorrhiza types besides the focal AM or EcM. This field also includes suggestions for mycorrhiza-like associations in rootless plants such as hepatics (levels ‘AM-like’, ‘EcM-like’, ‘ErM-like’ and ‘OrM-like’). The fields ‘AM_intensity’ and ‘AM_frequency’ indicate relative intensity (estimated part of root system) and frequency (% of plant individuals colonized), respectively. Analogous fields exist for EcM, ErM and OM. The field ‘AM_method’ enables 17 options for indicating the method and scale for determination of AM, whereas ‘EcM_method’, ‘ErM_method’ and ‘OM_method’ offer ten, seven and seven options, respectively.

The FungalRoot database contains six remarks fields. The remark_mycorrhiza_type represents notes on reported mycorrhiza type or colonization determination method. Four fields enable expert opinions about mycorrhizal type of each observation reported in the database. The fields contain name of the expert (two fields, for two experts respectively) and the expert comment. These categories warn users against mycorrhizal type mis-assignments, which are common in the literature (Brundrett, 2017; Tedersoo & Brundrett, 2017; Brundrett & Tedersoo, 2018), while allowing to store in the database the data reported by the original publication. The current version of the database contains remarks of two experts: Leho Tedersoo and Mark Brundrett. However, the dynamic set-up of our database allows data additions and editing, with a possibility to add new comments by external experts. The field ‘other_remarks’ provides additional information about methods, specific experimental treatments, etc. used in each particular paper. Ecological and evolutionary analyses may be sensitive to such data (Brundrett M, 2018).

In order to facilitate use of the data and to enable efficient update and versioning, the currently published version of the FungalRoot database is built using MySQL programming language and is integrated to the online analysis work-bench PlutoF (Abarenkov *et al.*, 2010b). This structure enables management and editing of multiple fields, custom search by any field, and third-party annotations such as comments or specification of missing details.

## Results

### FungalRoot structure

Our data is freely available for the scientific community, upon citation of this article. The most updated version of the FungalRoot database and the Recommended mycorrhizal status for plant genera can be searched and downloaded at https://plutof.ut.ee/#/study/view/31884 [available upon acceptance]. The PlutoF platform enables data management by adding observations, metadata and alternative interpretations about data reliability. We invite scientific community to provide comments on the mycorrhizal status of individual species and genera, using the PlutoF planform. The current version of the database and of “Recommended list…” is provided as supplementary material (Table S3 and S1, respectively).

For data input, there are two principal ways: i) using an upload file in spreadsheet format or ii) direct data insertion over the web platform, which is analogous to the UNITE database system (Abarenkov *et al.*, 2010a). Both the online data insertion and upload file contain the same data fields supplied with specific information. Some fields contain free text, whereas others enable a selection menu to secure consistent terminology. The scientific terminology follows generally MIMARKS standards that were supplemented with more detailed terms (such as mycorrhiza types, specific methods, etc.) that are not covered by these standards.

### Mycorrhizal data

In total, our database contains 36,303 observations for 14,870 plant species. A total of 19,893 observations included in the database are linked to geographical coordinates (Figure 1).

**Figure 1.**
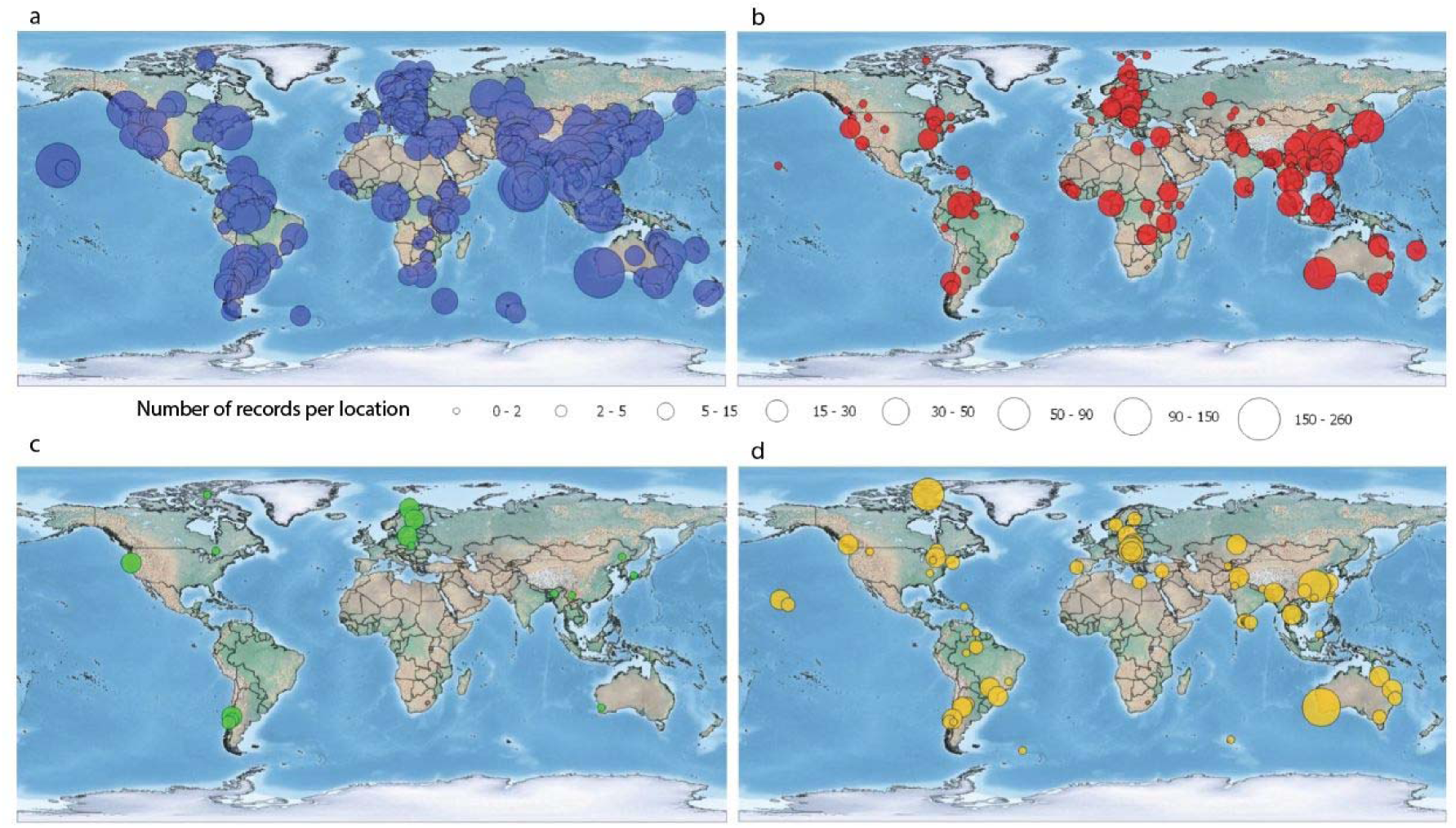
Georeferenced records included in the FungalRoot database. Circle size reflects number of observations per site. (a) arbuscular mycorrhizal colonization, (b) ectomycorrhizal colonization, (c) ericoid mycorrhizal colonization, (d) no mycorrhizal colonization.

Within the total number of observations, 45% and 2.5% include information about the intensity and frequency of mycorrhizal colonisation, respectively. Of mycorrhiza types, studies and observations about putatively AM plants prevail, followed by observations on EcM plants and non-mycorrhizal plants (Figure 2a). Among recorded habitats where mycorrhizal plants have been assessed, natural habitats prevail, being mostly forests and grasslands (Figures 2 b, c). Records are unequally distributed among plant species. Only 0.2% of the species had more than 40 records (Figure 3). Large number of species (59%) had only one record; 18 and 8% of species had 2 and 3 records respectively.

**Figure 2.**
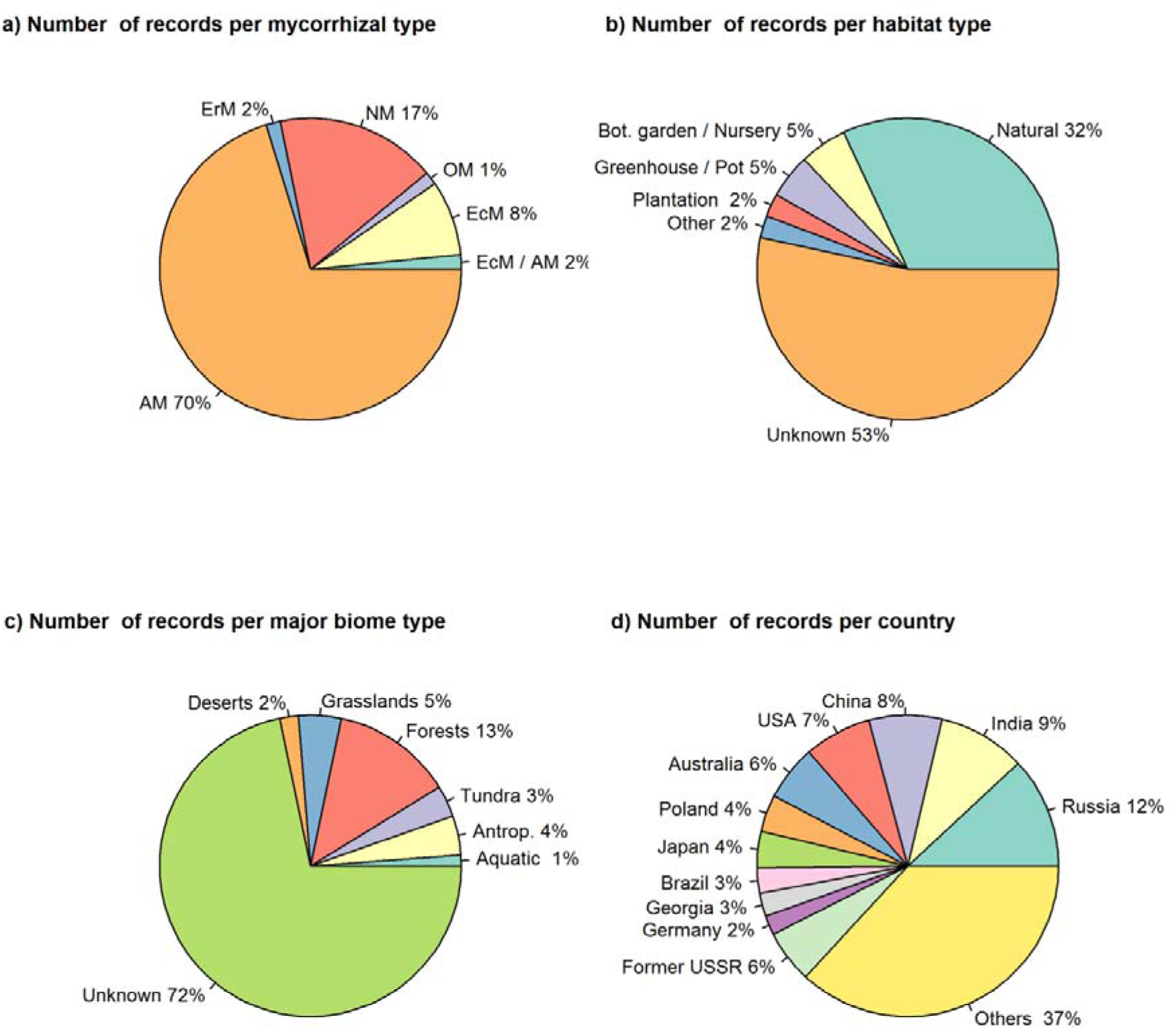
Number of records in the FungalRoot database (a) per most common mycorrhizal type, (b) per habitat type, (c) per major biome type, (c) per country. In the panel ‘a’ the EcM/AM category refers to the cases of mixed colonization by the two types of mycorrhizal fungi. The number of record for the types ‘ErM / AM’, ‘ErM / EcM’, ‘AM-like (non-vascular plants)’, ‘EcM-like (non-vascular plants)’, ‘ErM-like (non-vascular plants)’ and, ‘OM-like (non-vascular plants)’ is 9, 14, 8, 22, 0, 0, respectively. Due to small values these categories are not shown in the graph. In the panel ‘c’ the biome ‘Aquatic’ includes mangroves; The ‘Antrop.’ stays for ‘Atntropogenic’. In the panel ‘d’ the category ‘Former USSR’ refers to the records originated from the (Akhmetzhanova *et al.*, 2012) dataset, that are not assigned to Russia, but are assigned to other republics of USSR (now independent countries).

**Figure 3.**
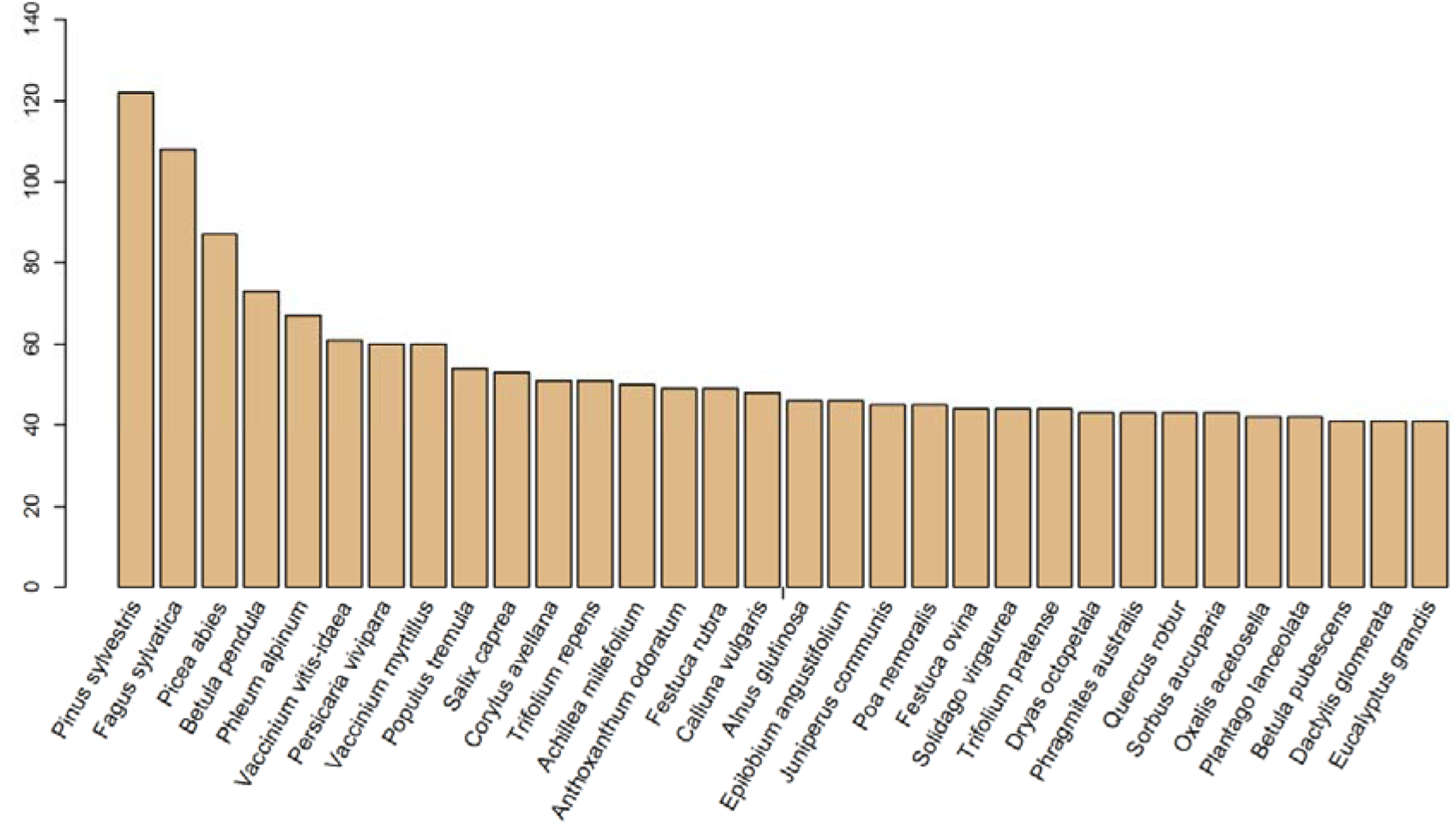
Plant species that have highest number of records (>40) in the FungalRoot database.

Observations about mycorrhizal status were unequally distributed globally, with greatest density in North Europe and North America and lowest density in Africa, Central Asia and Oceania (Figure 1). This is directly related to historical and present development of mycorrhiza research in different regions. We found literature about mycorrhizal status of plants in 9 languages that fit our criteria for inclusion. Relevant literature in English language clearly dominated, followed by Chinese, Spanish, Portuguese, Russian and French. Among the countries most of the plants has been examined in Russia, India, China and USA (Figure 2d).

In order to examine how distinct mycorrhizal types are distributed across plant growth forms (trees, herbs, shrubs), we extracted the publically available data from TRY (https://www.try-db.org/) (Kattge *et al.*, 2011). In this analysis, we considered the mycorrhizal type to correspond to that in the original report. to be AM/EcM/EcM all the plant species for which the respective mycorrhizal types are reported in the FungalRoot, summing up the records where only one mycorrhizal type is reported (i.e. all other types have been checked and not found) and the records simply reportinig the given mycorrhizal type. Among obligatory arbuscular mycorrhizal (AM, and EcM-AM plants) 50% are herbaceous, 25% are trees and the remaining plant species are distributed across the mycorrhizal types. Among facultatively arbuscular mycorrhizal (AM-NM) plants this ratio is 60/10/30. The great majority of ectomycorrhizal plants are trees and shrubs (92%) and the most of ericoid mycorrhizal plants are shrubs (85%). Among non-mycorrhizal plant species, 70% are herbaceous plants, 10% are trees and 20% belong to other growth forms (Figure 4).

**Figure 4.**
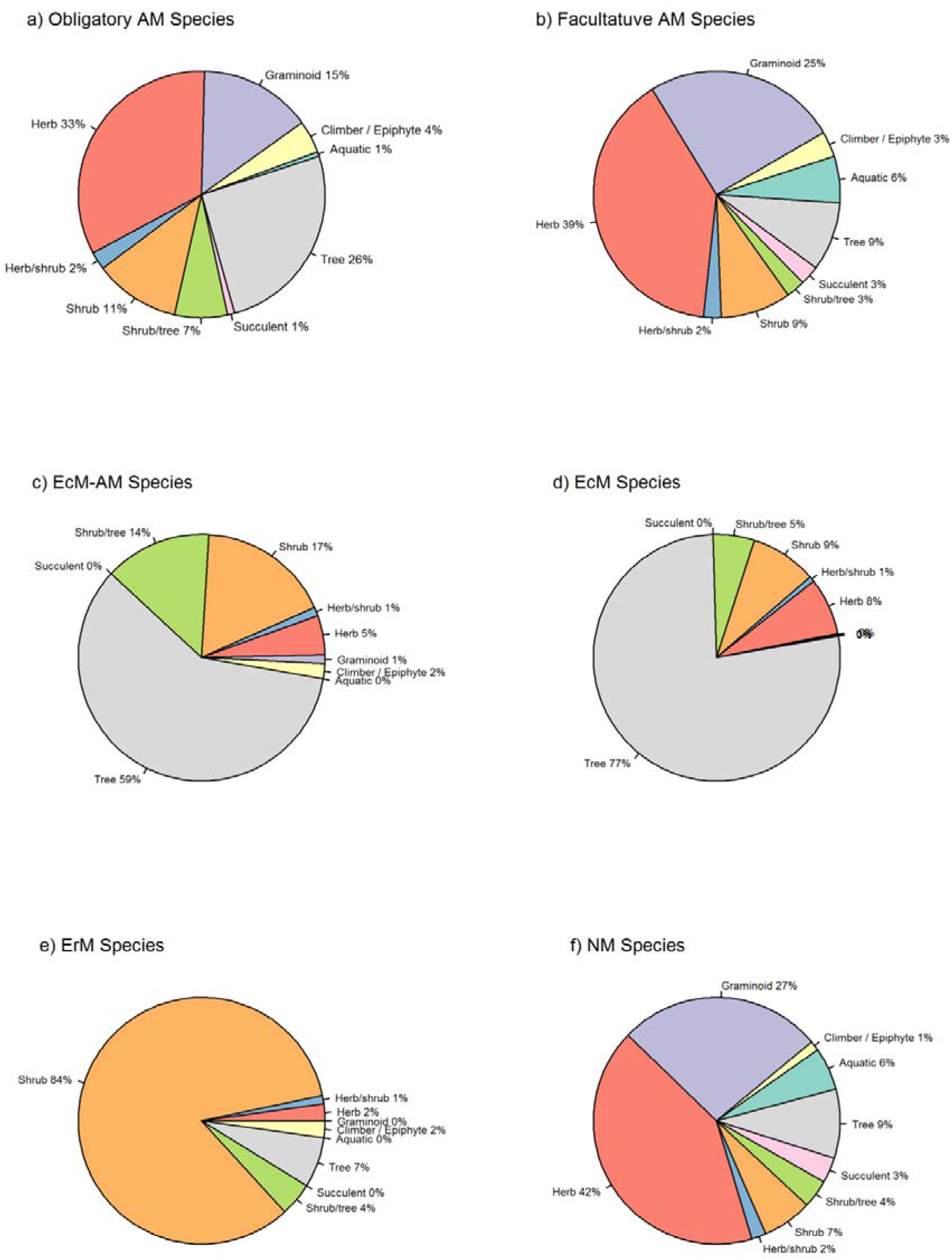
Distribution of plant growth form types across the main mycorrhizal types: AM – arbuscluar mycorrhizal plants, EcM ectomycorrhizal, ErM – ericoid mycorrhial, NM – non-mycorrhizal.

## Discussion

The FungalRoot database presented here provides species-by-site information about plant mycorrhizal associations and colonization intensity. We have significantly advanced previous attempts of such data compilations by exhaustive search for non-English literature, very old (>60 years) and recent literature, which resulted in tripling the number of species compared with the previously largest mycorrhizal type check lists of (Wang & Qiu, 2006), and (Akhmetzhanova *et al.*, 2012) that both contain records for ca 3000 plant species (overlapping to a large extent).

The database allows to summarize the contemporary information about the distribution of plant species per mycorrhizal type and distribution of mycorrhizal types per growth form. Our data confirms the earlier claims that the majority of mycorrhizal plants are arbuscular mycorrhizal (70% in our dataset), while despite wide ecological distribution (Read, 1991) ectomycorrhizal plants constitute only a tiny fraction of vascular plant species (0.7% in our dataset). However, given the fact that our data rather represent the research efforts in mycorrhizal studies than the true distribution of mycorrhizal plant species, these numbers should be treated with caution. Our data suggest that only ca 5% of all ca 400,000 vascular plant species have been examined for mycorrhizal type, with tropical plants being particularly understudied. Thus, further research is needed to obtain a truly quantitative understanding of patterns of mycorrhizal types distributions among vascular plants.

Despite the generally accepted view that the majority of EcM and ErM plants are shrubs and trees, while AM and not mycorrhizal habit are more or less equally distributed among plant growth forms, quantitative analyses on distribution of plant mycorrhizal types among growth forms has not been conducted till now. The data shown in the Figure 3 constitute the first attempt of quantitative exploration of thus far available information about mycorrhizal types of plant growth forms. The question what aspects of plant and mycorrhizal fungal physiology enable the overwhelming prevalence of woody forms among ectomycorrhizal and ericoid mycorrhizal plants is particularly intriguing. Further ecophysiological analyses of growth form preferences among plant mycorrhizal types will allow linking spatial patterns of plant functional types distributions to mycorrhizal habits. Given that the majority of ecological models of regional and global vegetation distribution and ecosystem functioning are based on plant functional types, this information will advance our understanding of impacts of mycorrhizas on functioning of terrestrial ecosystems.

Erroneous mycorrhizal diagnoses often provided in old literature and their blind, uncritical use has resulted in biased or incorrect interpretations of mycorrhizal type effects on evolutionary, biogeographic and ecophysiological processes (Brundrett & Tedersoo, 2019; Tedersoo *et al.*, 2019). To overcome these issues, we compared the original records with expert opinions derived from the rest of the data and other publications to construct a recommendation list for plant mycorrhizal associations (Table S1). It must be, however, noted that using this list uncritically has the following limitations: 1) it provides insufficient information about individual species and the effect of edaphic and climatic effects on mycorrhizal status; and 2) it may offer erroneous assignments to facultatively mycorrhizal taxa in ecosystems that are early successional, or exhibit extreme levels of nutrients or climatic conditions, such as alpine, flooded or fertilized habitats. In such cases, we recommend considering species-level assignments, provided in the FungalRoot database, accompanied by the edaphic data from specific regions or biomes, available as metadata in FungalRoot database. For species and genera not covered in FungalRoot database, we strongly recommend in situ determination of mycorrhizal types and mycorrhizal colonisation to reduce the determination biases.

In conclusion, the FungalRoot database features a number of unique characteristics, which will enrich the possibilities of scientific research based on the compiled metadata about locality, biome and edaphic conditions of the plant root sampling points. Such data enables quantitative analyses of drivers of mycorrhizal fungal colonization and distribution of mycorrhizal types, needed in order to understand the impacts of mycorrhizal symbiosis on functioning of the human-affected ecosystems. Furthermore, the database records have been traced to original publications, which enabled us to eliminate duplicated records caused by combining information from multiple compilations. The thorough quality check of the of mycorrhizal type data in the database, alongside with the recommendations for the genus-level mycorrhizal colonization type (Table S1) considerably reduce the amount of flaws in scientific studies addressing mycorrhizal type effects. Therefore, our database can be readily used for assessing the ecophysiological roles of mycorrhizal types in plant communities and ecosystem services and in comparative phylogenetics analyses targeting trait evolution. When coupled to other plant trait, ecological, evolutionary, soil and climate data, the FungalRoot database enables to test large-scale hypotheses about global processes such as biogeochemical nutrient cycling, climate change impact, and co-evolution of plants and fungi.

## Acknowledgements

LT received funding from the Estonian Science Foundation (PUT1399, MOBERC). NS, MB and SV were supported by the vidi grant 016.161.318 provided by the Netherlands Organization for scientific research (NWO), issued to NS.

